# Design Rules for Expanding PAM Compatibility in CRISPR-Cas9 from the VQR, VRER and EQR variants

**DOI:** 10.1101/2025.09.03.673821

**Authors:** Francisco Vieyra, Chinmai Pindi, George P. Lisi, Uriel N. Morzan, Giulia Palermo

## Abstract

Expanding the range of Protospacer Adjacent Motifs (PAMs) recognized by CRISPR-Cas9 is essential for broadening genome-editing applications. Here, we combine molecular dynamics simulations with graph-theory and centrality analyses to dissect the principles of PAM recognition in three Cas9 variants – VQR, VRER, and EQR – that target non-canonical PAMs. We show that efficient recognition is not dictated solely by direct contacts between PAM-interacting residues and DNA, but also by a distal network that stabilizes the PAM-binding domain and preserves long-range communication with REC3, a hub that relays signals to the HNH nuclease. A key role emerges for the D1135V/E substitution, which enables stable DNA binding by K1107 and preserves key DNA phosphate locking interactions via S1109, securing stable PAM engagement. In contrast, variants carrying only R-to-Q substitutions at PAM-contacting residues, though predicted to enhance adenine recognition, destabilize the PAM-binding cleft, perturb REC3 dynamics, and disrupt allosteric coupling to HNH. Together, these findings establish that PAM recognition requires local stabilization, distal coupling, and entropic tuning, rather than a simple consequence of base-specific contacts. This framework provides guiding principles for engineering Cas9 variants with expanded PAM compatibility and improved editing efficiency.

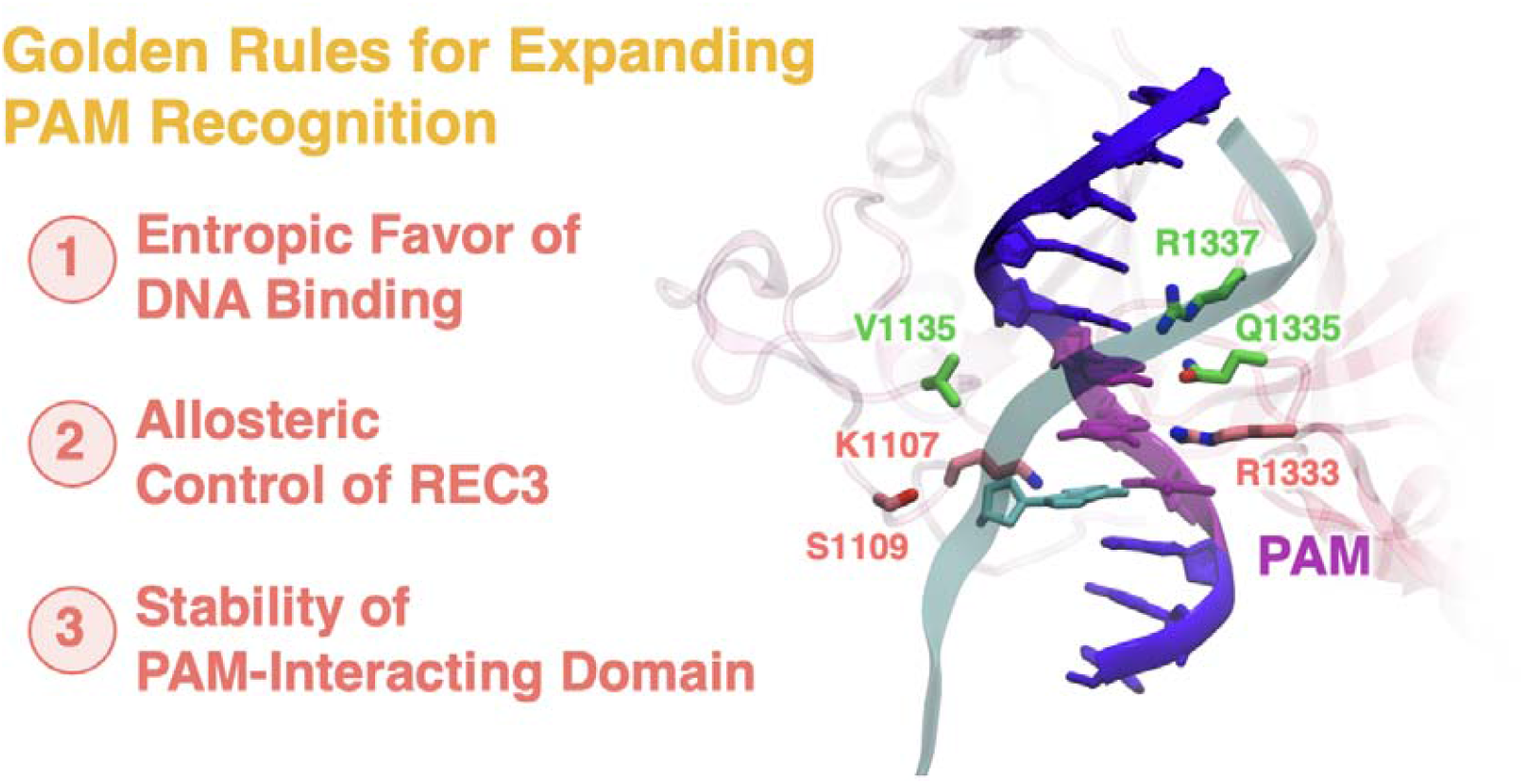

## Introduction

Expanding the DNA sequence recognition capabilities of CRISPR-Cas9 is a key priority for enhancing the versatility and precision of this powerful genome editing system^1^. While CRISPR-Cas9 has revolutionized the life sciences by enabling facile and programmable genome editing^2^, broadening its targeting scope remains essential for achieving greater precision across diverse biotechnological applications^1^.

In this system, the Cas9 endonuclease complexes with a guide RNA to recognize and cleave complementary DNA sequences^3^. Target recognition is initiated by the binding of a Protospacer Adjacent Motif (PAM) to a groove formed by Cas9’s C-terminal region and a domain structurally related to type II Topoisomerase (**Figure 1a**)^4^. This PAM interaction enables the guide RNA to hybridize with the target DNA strand (TS), displacing the non-target strand (NTS). Cas9 is a large, multi-domain protein, with RNA binding mediated by an α-helical recognition (REC) lobe. DNA cleavage is catalyzed by two nuclease domains: HNH, which cleaves the TS, and RuvC, which cleaves the NTS, resulting in a double-strand break.

**Figure 1.**
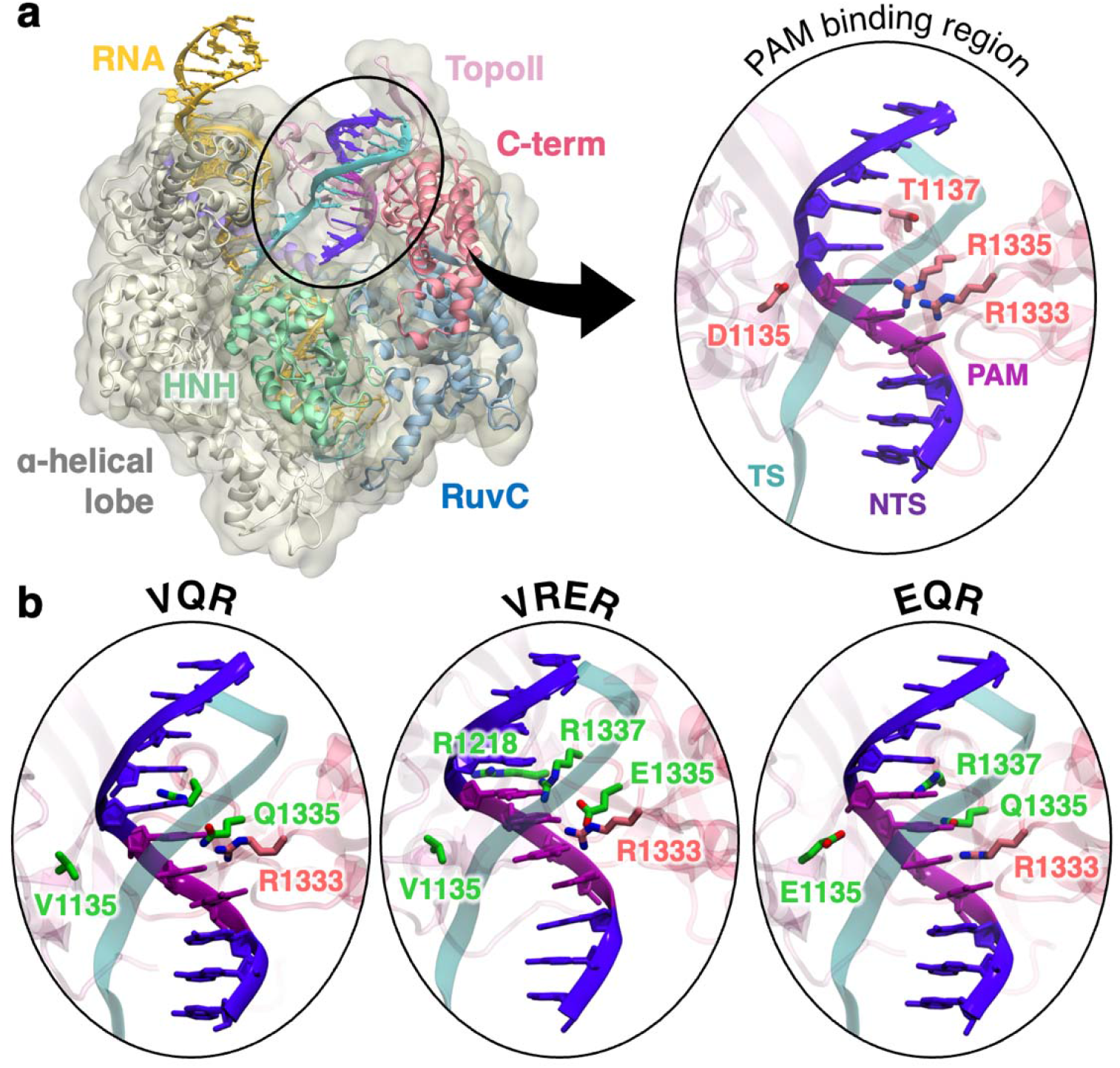
Overview of the *S. pyogenes* CRISPR-Cas9 system and its VQR, VRER and EQR variants. **a**. Architecture of CRISPR-Cas9 and detailed view of the PAM recognition region (PDB: 4UN3)^4^. Cas9 is shown in molecular surface, highlighting the protein domains with colored ribbons. The RNA (yellow) and the DNA target strand (TS cyan) and non-target strand (NTS, violet) are also shown as ribbons. The PAM sequence is shown in magenta. **b**. VQR (PDB: 5FW1), VRER (PDB: 5FW3) and EQR (PDB: 5FW2)^11^ variants of CRISPR-Cas9 with mutations in the PAM recognition region.

The binding of the ‘5-TGG-3’ PAM sequence to the R1333 and R1135 residues is a critical prerequisite for DNA recognition and subsequent cleavage by Cas9^2,5^. This strict PAM requirement imposes a major constraint on the breadth of Cas9-mediated genome editing^1^. Indeed, while the guide RNA can be readily reprogrammed to target virtually any DNA sequence, the effective recognition and cleavage remain restricted to targets adjacent to the ‘5-TGG-3’ sequence. To overcome this limitation and expand the range of targetable PAM sequences, Cas9 has been engineered via directed evolution to exhibit altered PAM specificities^6^. Notably, three early variants of Cas9 have been developed (**Figure 1b**): the VQR variant (D1135V, R1335Q, T1337R), which recognizes 5’-NGA-3’; the VRER variant (D1135V, G1218R, R1335E, T1337R), which recognizes 5’-NGCG-3’; and the EQR variant (D1135E, R1335Q, T1337R), which recognizes 5’-NGAG-3’. These variants retain robust genome-editing activity while accommodating alternative PAM sequences, thus broadening the targeting scope of the CRISPR-Cas9 system. In contrast, variants carrying only substitutions of the arginine residues that directly contact the canonical 5′-TGG-3′ PAM (i.e., R1335/R1335 ⍰ Q/E) abolish DNA cleavage activity without altering PAM specificity^6^. This observation underscores the cooperative nature of PAM recognition and catalysis^7,8^ and is intriguing. Indeed, substituting arginine (R) with glutamine (Q) is expected to enhance recognition of adenine-containing PAMs – given glutamine’s affinity for adenine – whereas native arginine preferentially interacts with guanine ^9,10^.

This suggests that while mutations in PAM-interacting residues are essential, they are not sufficient to reprogram PAM specificity. Additional substitutions, such as D1135V, are required for effective PAM reprogramming, yet their functional roles remain unclear. Notably, D1135V is located distal to PAM and does not contact the DNA or the PAM-binding residues (**Figure 1b**)^11^, raising the question of how this mutation contributes to altered PAM specificity. While the plasticity of the PAM-binding cleft may facilitate adaptation to different PAM sequences^12,13^, the specificity and underlying mechanism is not understood.

Here, we performed molecular dynamics (MD) simulations on three Cas9 variants – VQR, VRER, and EQR – alongside wild-type Cas9 (wt-Cas9) and two engineered constructs carrying R1333Q/R1335Q substitutions that fail to alter PAM specificity. By integrating graph-theoretical and centrality analyses, we show that efficient PAM recognition emerges from three interdependent features: stabilization of the PAM-interacting domain, preservation of long-range allosteric communication with the REC3 hub, and entropic tuning of DNA engagement. These principles highlight that the PAM-interacting domain functions not merely as a local recognition module but as an upstream allosteric hub, coupling PAM sensing to distal conformational changes required for HNH activation. Our findings therefore underscore that engineering Cas9 variants with altered specificity requires consideration of distal allosteric networks and global communication pathways, not just immediate base-specific contacts at the PAM site.

## Computational Methods

### Molecular Dynamics (MD) Simulations

MD simulations were carried out on the wild-type *S. pyogenes* Cas9 (wt-Cas9) and its VQR, VRER and EQR variants, based on the X-ray structures from the Protein Data Bank: 4UN3 for wt-Cas9 (2.58 Å)^4^, 5FW1 for VQR (2.65 Å), 5FW3 for VRER (2.77 Å), and 5FW2 for EQR (2.75 Å)^11^. These structures represent comparable conformational states of the complex. Two systems that fail to expand PAM recognition were also examined: one featuring the R1333Q and R1335Q mutations within the PAM-interacting domain (*viz*., ENG1), and a second in which the PAM sequence was altered from the canonical 5’-TGG-3’ to 5’-TAA-3’ (*viz*., ENG2). Each model system was embedded in explicit water, resulting in periodic simulation boxes of ∼180*116*139 Å^3^, containing ∼270K atoms. MD simulations were performed using a protocol tailored for protein/nucleic acid complexes^14^, also applied in recent studies of CRISPR-Cas genome editing systems^15–18^. The Amber ff19SB^19^ force field was employed, incorporating the OL15^20^ and OL3^21^ corrections for DNA and RNA, respectively. The TIP3P model was used for water^22^. Simulations were performed in the NPT ensemble with temperature held at 310 K using the Bussi thermostat^23^. The pressure was held at 1 bar with the Parrinello-Rahman barostat^24^. The Particle Mesh Ewald method was used to compute long-range electrostatic interactions with a cut-off radius of 1 nm. Bonds involving hydrogen were restrained using LINCS. The equation of motion was integrated using a time step of 2 fs. For each system ∼1 µs of MD simulations were carried out in three replicates. All simulations were performed using Gromacs (v. 2024.0)^25^.

### Community Network Analysis

To characterize the allosteric network and how it is altered in the VQR, VRER and EQR variants, and in the ENG1 and ENG2 systems, we performed Community Network Analysis (CNA)^26^. Here, the system is described as a network of node residues connected by edges whose lengths inversely reflect motion correlations, according to *w*_*ij*_ *=* − log *CG*_*ij*_ (where *GC* are the Generalized Correlations)^27^, thereby generating molecular networks for Cas9 and its variants^28^. These correlations were derived from Shannon entropy to capture both linear and non-linear coupling. Dynamical network models were then divided in local substructures – i.e., communities – composed of groups of nodes in which the network connections were dense but between which they were sparse. To structure the communities, the Girvan-Newman graph-partitioning approach^29^ was employed, using the Edge Betweenness (EB) as partitioning criterion (details are reported in the Supporting Information, SI). EB was defined as the number of shortest pathways that cross the edge, thereby accounting for the number of times an edge acts as a bridge in the communication flow between nodes of the network. The total EB between couples of communities (i.e., the sum of the EB of all edges connecting two communities) is a measure of their communication strength.

### Communication Pathway Analysis

To map allosteric communication pathways, we applied Dijkstra’s shortest-path algorithm to our molecular networks^30^. The resulting pathways consist of sequential edges that maximize total correlation – thereby optimizing momentum transport – between designated “source” and “sink” residues. For this study, we computed communication pathways between the DNA recognition site in the REC3 domain and the DNA cleavage site. Specifically, α-helix 37 (residues 692–699) in REC3 was defined as the source, and the HNH catalytic residue H840 as the sink, reflecting the established importance of REC3– HNH allosteric communication for Cas9 function and specificity^31,32^. To evaluate the impact of auxiliary signaling routes, we identified the 50 highest‐scoring paths for each source–sink pair, normalized their contributions, and mapped the aggregated network onto the 3D structures of wt-Cas9 and its variants. This approach captures the influence of alternative, suboptimal pathways that may complement or compete with the optimal route, thereby providing a more comprehensive view of the allosteric communication landscape^33–39^. In our recent studies^32,40–42^, this approach uncovered both the primary allosteric channels and secondary pathways that bolster signal robustness, offering a comprehensive view of dynamic signal propagation through the system.

#### Eigenvector Centrality Analysis

Eigenvector Centrality (EC) is a centrality measure, which quantifies the influence of a node by weighting it according to the importance of its neighbors within the network’s dynamics^28,43^. The EC of a node, *c*_*i*_, defined as the sum of the centralities of all nodes that are connected to it by an edge, is 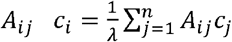, where the edges *A*_*ij*_ are elements of the adjacency matrix *A* (based on Generalized Correlations), and *λ* is the eigenvalue associated to the eigenvector composed by *c*_*i*_ elements.

This approach assigns the system’s functional dynamics to its dominant collective mode (the first eigenvector of *A*). Accordingly, *λ* corresponds to the largest eigenvalue of *A*, which –by the Perron–Frobenius theorem – is guaranteed to be real, positive, and unique. The EC estimation quantifies the degree of connectivity of each node within the system, therefore measuring how well the nodes of the protein network are connected to other well-connected nodes. At variance with other centrality descriptors^33^, the EC serves as a measure of the connectivity against a fixed scale when normalized and can reliably compare the system’s connectivity associated with mutations or effector binding, as applied here for the analysis of the wt-Cas9 and its variants.

## Results

### Dynamic Remodeling of Cas9 Variants

To gain molecular insights into PAM selectivity, we conducted MD simulations of the VQR, VRER, and EQR variants, along with two engineered systems. The first (ENG1) features glutamine substitutions at residues R1333 and R1335 (R1333Q/R1335Q) that are expected to weaken guanine interactions while enhancing affinity for adenine^9,10^. To reflect this altered specificity, we also introduced a 5’-TAA-3’ PAM sequence in a second system (ENG2).

To assess how different mutations influence the dynamics and inter-domain communication within Cas9, we performed community network analysis (CNA) across all variants (**Figure 2**)^26^. CNA identifies groups of highly correlated residues – referred to as communities and quantifies the strength of communication between them based on the edge betweenness metric (see Methods). In the resulting network, inter-community connections are represented as edges, whose thickness reflect the degree of information flow. This approach is particularly well-suited for characterizing the protein’s allosteric architecture and reveals how communication pathways are reshaped by specific mutations.

**Figure 2.**
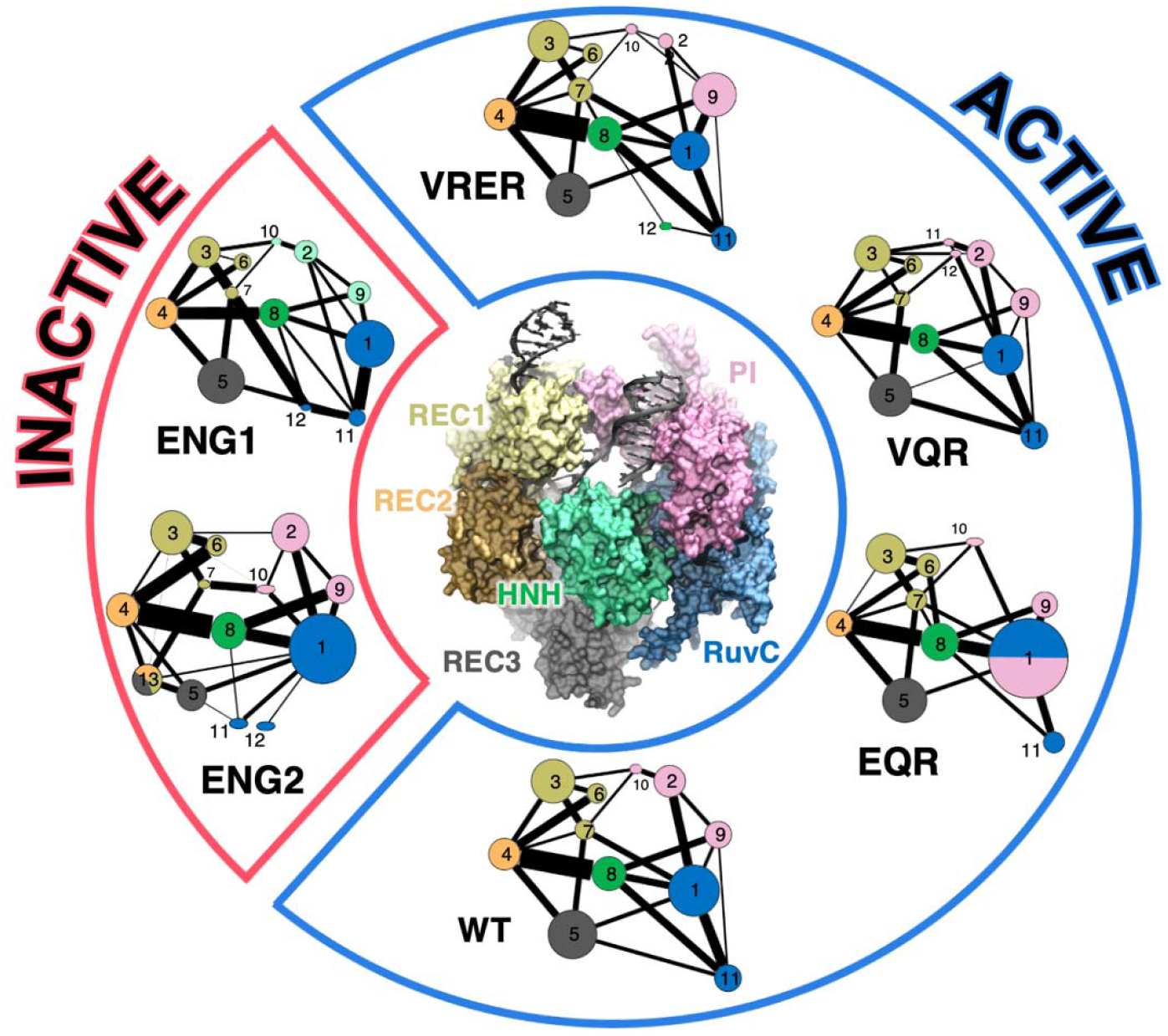
Community structure of CRISPR-Cas9 and engineered variants. The network diagrams represent the community architecture of the wt-Cas9, active variants (EQR, VQR, VRER), and inactive engineered variants (ENG1, ENG2). Each community corresponds to a protein domain, while edges represent the strength of inter-domain communication based on dynamic correlations. Thicker edges indicate stronger coupling. The central structure shows the domain organization of Cas9, with HNH (green), RuvC (blue), PI (pink), REC1–3 (shades of gray to yellow). Active variants exhibit tighter and more integrated community networks compared to the fragmented architecture observed in inactive forms.

The community distribution in wt-Cas9 and its active variants closely mirrors the underlying domain architecture (**Figure 2**), consistent with our earlier CNA analysis of wt-Cas9 dynamics^8^. In contrast, the inactive variants (ENG1 and ENG2) exhibit a more fragmented community organization in REC3 and RuvC, a characteristic feature of the previously reported PAM-unbound state^8^. Notably, in all active variants, the strongest edge in the network connects the HNH and REC2 domains, reflecting a high degree of dynamical correlation between these regions. This observation is consistent with previous studies demonstrating that REC2 plays a critical role in modulating HNH catalytic activity^31,32,44,45^. Both computational and single-molecule analyses have shown that REC2 and HNH undergo coordinated conformational changes to enable proper DNA binding and cleavage. In contrast, this interdomain edge is markedly weakened in the engineered variants, suggesting that even subtle perturbations in the PAM-interacting region can disrupt the long-range communication required for HNH activation.

To further assess allosteric signaling in the studied variants and wt-Cas9, we applied Eigenvector Centrality (EC) analysis (**Figure 3**)^43^. This metric evaluates the relative importance of each node within the network, with EC specifically capturing a node’s influence by weighting its connections according to their contribution to the system’s dynamics (see Methods). When applied to networks derived from MD simulations, EC highlights residues and regions that drive global collective dynamics, making it a powerful tool to pinpoint essential amino acids involved in long-range allosteric communication.

**Figure 3.**
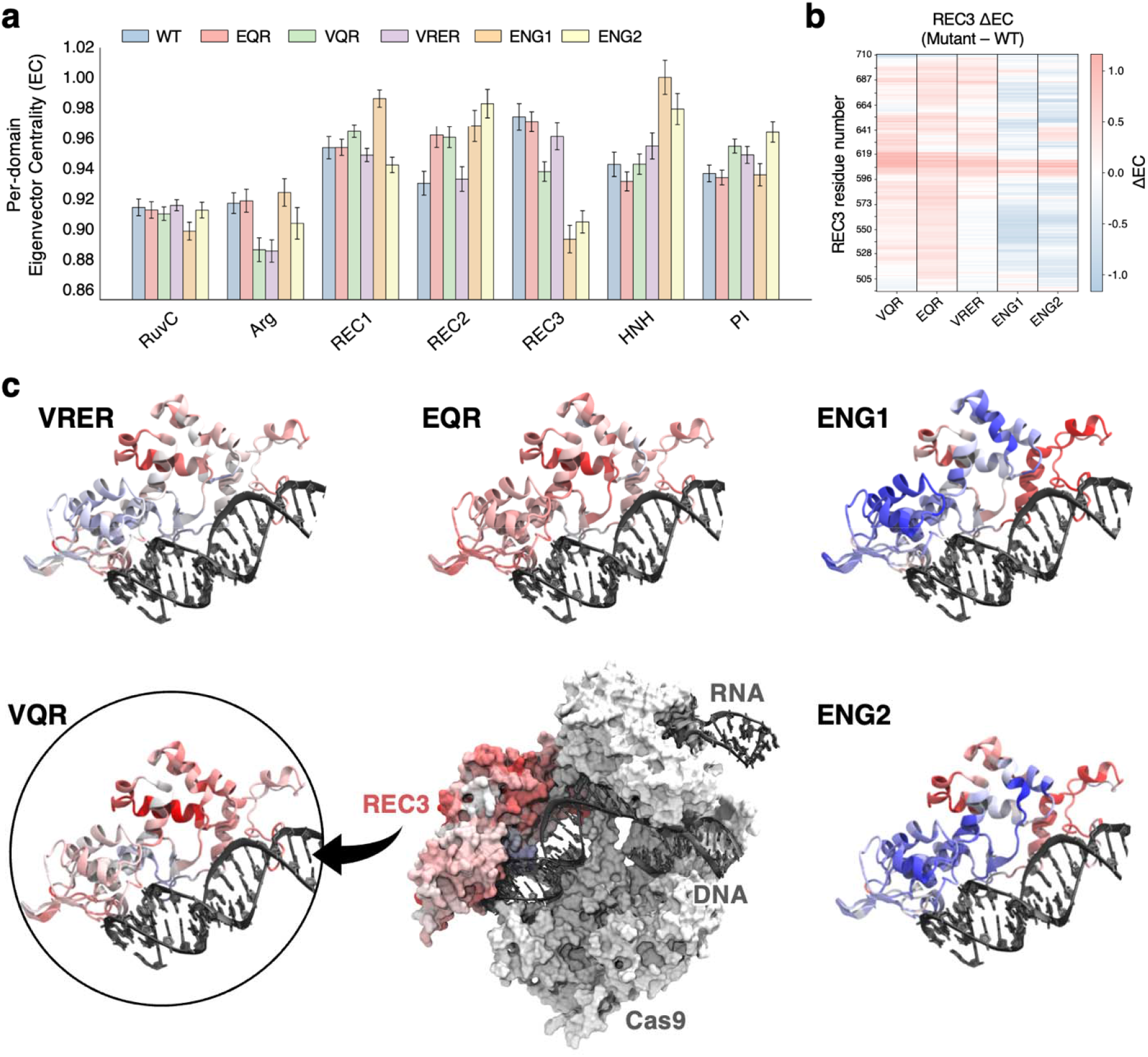
Eigenvector Centrality analysis reveals global allosteric remodeling across Cas9 variants. **a**. Per-domain Eigenvector Centrality (EC) values show reduced dynamic centrality in the REC3 domain for inactive variants (ENG1, ENG2), suggesting altered allosteric communication linked to disrupted PAM interactions. Values are averaged over three independent simulation replicates and normalized over the number of residues. Error bars represent the standard deviation of the mean, computed using block averaging. **b**. Heatmap of mutation-induced EC differences (variant − wt) in the REC3 domain shows pronounced dynamic shifts in the ENG1 and ENG2 variants. **c**. Structural mapping of EC differences (variant − wt) onto Cas9–DNA complexes, highlighting dynamical alterations in the REC3 domain of the inactive engineered variants, distinguishing them from the active forms.

The EC profiles quantified the centrality of each residue within the protein network (**Figure S1**). Analysis of the domain-averaged EC values (**Figure 3a**) consistently shows that the HNH domain exhibits higher centrality than all other domains across systems, highlighting it as the principal hub of Cas9 dynamics. This observation aligns with previous studies highlighting the intrinsic flexibility of HNH and its essential role in CRISPR-Cas9 function^7,46^, where conformational changes in this domain are key to enabling catalysis. Interestingly, the inactive variants (ENG1 and ENG2) show a pronounced reduction in centrality within the REC3 domain.

This suggests that mutations limited to the PAM-interacting residues R1333 and R1335 perturb distal REC3 sites through long-range allosteric communication.

The contrasting REC3 centrality between active and inactive variants underscores that tightly coordinated REC3 dynamics are essential for Cas9 function, and that disruption of this synchronization can abolish activity. It is also notable that the variability of the EC values in the REC3 domain – reflected by the increased fluctuations in the per-residue EC (**Figure S1**) – is higher in the ENG1 and ENG2 inactive variants compared to the active ones. The increased fluctuations in REC3 centrality suggest a more heterogeneous dynamic behavior in the inactive variants, reinforcing the idea of a dynamic coupling between the PAM-interacting domain and REC3^12^. Taken together, these findings demonstrate that mutations near the PAM-binding site can induce significant global changes in the dynamic architecture of CRISPR-Cas9.

### Effect of Mutations on PAM Recognition

Here, we analyze the local dynamics and interactions within the PAM-interacting domain of Cas9 and its variants. This analysis provides insight into how specific mutations locally modulate DNA binding. In combination with our global CNA and EC analyses (*vide supra*), this links local structural perturbations at the PAM-binding site to broader changes in the protein’s dynamic and allosteric architecture.

In the VQR and VRER systems, the PAM-interacting residues R1333 and Q1335 (glutamate in VRER) consistently maintain stable interactions with the PAM sequence (**Figure 4a-c**). Additionally, R1337 (mutated from threonine in wt-Cas9) engages with the third PAM guanine in VQR, and with the phosphate backbone of the third PAM cytosine in VRER (**Figure 4a, d**), further stabilizing the protein–DNA interface. These interactions are lost in the engineered variants, where R1333 and R1335 are mutated to glutamines (**Figure 4a-d**). The disruption is particularly evident in the ENG2 system, which features glutamine substitutions alongside an adenine-rich PAM. Despite the reported preference of glutamine for adenine, these mutations fail to establish stable contacts, suggesting that modifying PAM-interacting residues alone is insufficient to ensure PAM recognition.

**Figure 4.**
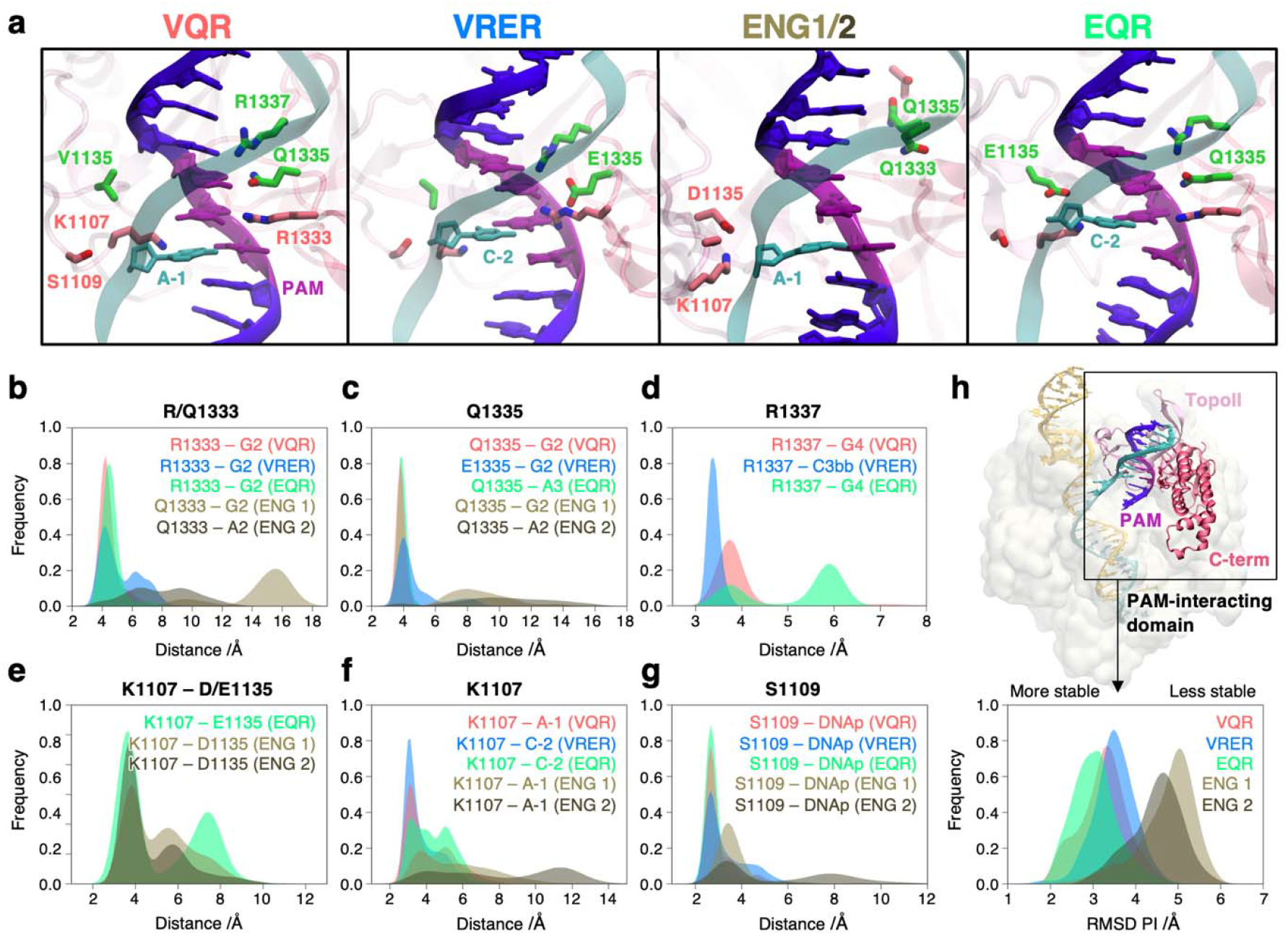
Local interactions and conformational stability within the PAM-interacting domain of Cas9 variants. **a**. Representative snapshots highlight key PAM-contacting residues in VQR, VRER, EQR, and engineered variants (ENG1/2), showing that R1333, Q/E1335, and R1337 maintain stable interactions with the PAM in active variants but not in engineered forms. **b-g**. Distributions of key residue–residue and residue–DNA distances that modulate PAM recognition and DNA engagement. **b**. R/Q1333 and PAM bases; **c**. Q/E1335 and PAM bases; **d**. R1337 and DNA; **e**. D/E1135 and K1107, highlighting the formation of a stabilizing salt bridge; **f**. K1107 and DNA bases, showing DNA engagement in the active Cas9 variants; **g**. S1109 and the DNA phosphate backbone, illustrating the stability of the “phosphate lock” interaction in the active variants. **h**. Root mean square deviation (RMSD) of the PAM-interacting domain, showing diminished structural stability in the ENG1/2 variants relative to VQR, VRER, and EQR. The RMSD is computed for the backbone atoms with respect to the equilibrated structures. The PAM-interacting domain includes the C-terminal and the type II Topoisomerase–like (TopoII) domain.

To further investigate this observation, we analyzed the interaction network of D1135, a key residue mutated to valine in the VQR and VRER variants, and to glutamate in EQR. Although this mutation is conserved across active variants, its functional role remains poorly understood. Indeed, this residue is positioned on the opposite side of the DNA double helix, away from the PAM-interacting interface. In the ENG1-2 systems, D1135 forms a stable salt bridge with K1007 (**Figure 4a, e**), an interaction that is absent in VQR and VRER due to the D1135V mutation. Consequently, K1007 is free to interact persistently with DNA bases in VQR and VRER (**Figure 4f**). In contrast, the D1135–K1007 salt bridge in the engineered variants sequesters K1107 from DNA, destabilizing its interaction. This destabilization is further reflected by the loss of the critical “phosphate lock” interaction between S1109 and the DNA phosphate backbone (**Figure 4g**), essential for substrate stabilization. Notably, in the inactive engineered variants, the loss of PAM-DNA interactions combined with the repositioning of key residues on the opposite side of the DNA helix leads to a marked destabilization of the PAM-interacting domain compared to the active VQR, VRER, and EQR variants (**Figure 4h**). Interestingly, in the EQR variant, D1135 is replaced by glutamate, which can potentially preserve the salt bridge with K1107. This interaction is transiently maintained during the simulation but does not prevent K1107 from interacting with the DNA bases (**Figure 4e-f**). Hence, in the EQR variant, the D1135E substitution does not sequester K1007 from DNA, but rather contributes to its stable location in proximity of DNA. Previous studies have also shown that single-point mutations such as E1219V remodel the hydration and electrostatic environment of the PAM pocket^47^.

Consistent with this, our results reveal that additional distal substitutions (D1135V/E) are essential to stabilize the broader interaction network required for effective PAM recognition.

Overall, these findings suggest that the D1135V mutation is critical for stabilizing the DNA duplex in the VQR and VRER variants, by enabling K1107 to engage directly with the DNA. This interaction, in turn, enables the shorter glutamine side chain at position 1335 (mutated from the longer arginine in wt-Cas9) to effectively contact the PAM bases. In the EQR variant, the D1135E substitution does not sequester K1107 from the DNA; rather, it appears to support complex stabilization by maintaining both intra-protein interactions and productive DNA contacts.

Together, these results demonstrate that PAM recognition in Cas9 is not determined solely by direct contacts between PAM-interacting residues and the DNA but also depends on a network of distal interactions that stabilize the PAM-interacting domain. Disruption of this network, as observed in the inactive ENG1/2 variants, compromises DNA binding at the PAM site.

### Alteration of the HNH–REC3 Allosteric Signaling

Cas9 is an allosteric enzyme in which DNA binding triggers activation of distal nuclease sites^48,49^. A key allosteric pathway links the HNH nuclease to the REC3 domain. Single-molecule experiments have shown that mutations within REC3 – such as those in high-fidelity Cas9 variants designed to reduce off-target effects^50^ – modulate HNH dynamics and, in turn, its activation for DNA cleavage^31^. This HNH–REC3 communication has therefore been proposed as a critical determinant of the mechanism underlying the enhanced specificity of these variants.

Using a combination of NMR, MD simulations and graph theory, we have recently shown that this communication is mediated by α-helix 37 (residues 692–699)^32^, which senses target DNA complementarity^51^. We found that high-specificity mutations in REC3 increase the communication between α-helix 37 and HNH, providing a structural basis for the enhanced specificity variants.

Here, to investigate how alterations in the PAM-binding site influence HNH–REC3 communication, we computed allosteric pathways between α-helix 37 and the catalytic residue H840 using the Dijkstra algorithm. This analysis identifies shortest paths that maximize total correlation, thereby minimizing the communication distance between these spatially distinct sites (see Methods). In addition to the optimal pathway, the 50 most probable suboptimal information channels were computed, aggregated, and mapped onto the 3D structure (**Figure 5**), thereby accounting for the contribution of alternative communication routes.

**Figure 5.**
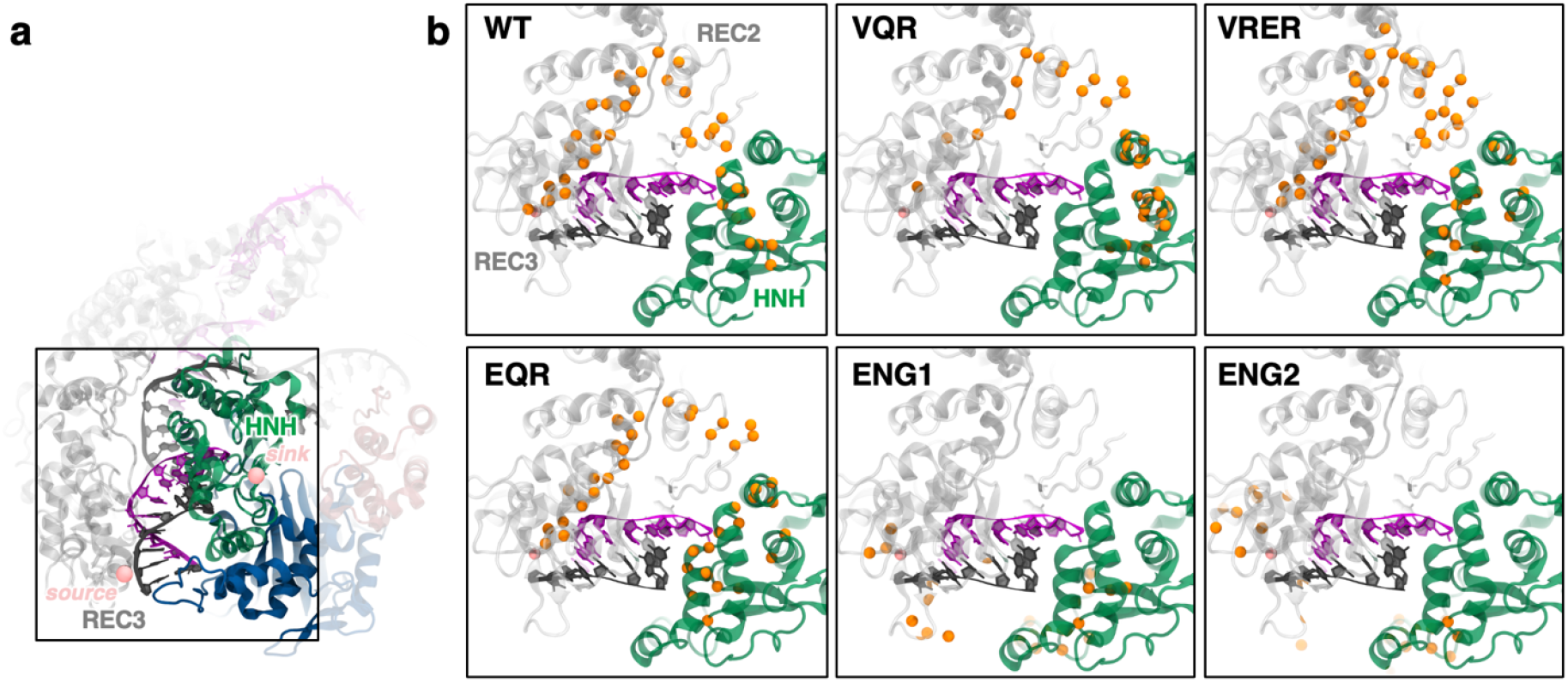
Allosteric communication pathways from REC3 to HNH. **a**. Overview of Cas9 highlighting the REC3 and HNH domains involved in long-range communication. **b**. Allosteric pathways from α-helix 37 (residues 692–699, source) to the HNH catalytic residue H840 (sink) were computed using Dijkstra’s shortest-path algorithm for each Cas9 variant. The top 50 highest-scoring suboptimal pathways are shown as red spheres mapped onto the 3D structure. In wt-Cas9 and the active VQR, VRER, and EQR variants, a continuous communication pathway from REC3 to HNH is preserved. This pathway is disrupted in the inactive engineered variants ENG1 and ENG2, reflecting impaired long-range communication due to destabilization of the PAM-interacting domain (**Figure 4h**).

In wt-Cas9 and the active VQR, VRER, and EQR variants, a robust communication pathway between α-helix 37 and HNH is maintained. In contrast, this communication is disrupted in the ENG1 and ENG2 variants. In these inactive systems, the loss of key locking interactions between K1107 and DNA leads to pronounced destabilization of the PAM-interacting domain (**Figure 5b**). Consistently, the ENG1 and ENG2 variants display a pronounced decrease in REC3 domain centrality compared to the wild-type and active variants, reflecting a loss of coordinated dynamics. Together, these findings demonstrate that perturbations to the stability and dynamics of the PAM-interacting domain propagate to REC3, thereby disrupting the critical HNH–REC3 allosteric communication.

These results suggest a model in which the PAM-interacting domain serves as an upstream regulatory hub that modulates the conformational coupling between REC3 and HNH. Disruption of this hub – whether by loss of PAM–DNA contacts or destabilization of key interdomain interactions – compromises the integrity of the HNH–REC3 allosteric pathway, thereby impairing the HNH activation.

## Discussion

Here, we combined MD simulations with graph-theory–based network and centrality analyses to dissect the mechanistic basis of the expanded DNA recognition in CRISPR-Cas9.

Our results show that PAM recognition is governed not only by direct contacts between canonical PAM-interacting residues and DNA, but also by a broader network of distal interactions that stabilize the PAM-interacting domain and preserve long-range allosteric pathways. In the VQR, VRER, and EQR variants, this network remains intact, enabling coordinated coupling between the PAM-interacting domain, REC3, and HNH. In these systems, PAM-interacting residues form stable contacts with the noncanonical PAM, while the surrounding network positions K1107 to engage the DNA and S1109 to maintain the “phosphate lock”^4^, both essential for stable DNA binding (**Figure 4**). The observation that the distal D1135V mutation rescues the destabilization introduced by R1333/R1335 substitutions is consistent with thermodynamic analyses of Cas9-NG, a variant that broadens PAM compatibility from NGG to NGN sequences^52^, where the destabilizing effect of R1335V is mitigated by compensatory mutations such as E1219F and D1135V ^53^. By contrast, ENG1 and ENG2 lose persistent PAM contacts and form a D1135–K1107 salt bridge that sequesters K1107 from the DNA phosphate backbone, disrupts phosphate-lock stability, and increases structural fluctuations within the PAM-interacting domain (**Figure 4h**). This local destabilization propagates through the structure, disrupting REC3 dynamics (**Figure 3**) and breaking the HNH–REC3 communication pathway (**Figure 5**).

Community Network Analysis (CNA) illustrates how this local destabilization results in a fragmented REC3 and RuvC community architectures in the engineered inactive systems (**Figure 2**). Centrality analysis further illustrates how perturbations in the PAM-interacting domain propagate through and reshape the global dynamics of Cas9 (**Figure 3**). Inactive variants display a marked reduction in Eigenvector Centrality (EC) within REC3, accompanied by increased variance, indicating a more heterogeneous and less coordinated dynamic ensemble. These changes suggest that in ENG1 and ENG2, REC3 no longer operates as stable and efficient conduit for allosteric signaling. Mapping of the allosteric pathways between HNH and REC3 reinforces these findings. Using shortest-path analysis, we identified the highest-scoring communication routes linking the HNH catalytic core to α-helix 37 of REC3, a key structural element involved in sensing target DNA complementarity (**Figure 5**)^51^. The VQR, VRER and EQR variants preserve the communication between the key α-helix 37 and HNH. Whereas, in ENG1-2, the allosteric channels are markedly attenuated or rerouted, leaving α-helix 37 less effectively coupled to HNH. This disruption offers a direct molecular explanation for how local destabilization of the PAM-interacting domain propagates through the allosteric network to impair the allosteric activation of HNH.

Collectively, our findings support a model in which the PAM-interacting domain serves as an upstream allosteric hub, wherein structural stabilization of the PAM-binding site is essential for transmitting local DNA-binding events into long-range dynamical changes required for Cas9 function. This structural integrity is critical for maintaining the coupling between REC3 and HNH, ensuring effective allosteric communication. Mutations within the PAM-interacting domain that fail to preserve this local stability are insufficient to reprogram PAM recognition, underscoring that a stably anchored PAM-interacting domain is a prerequisite for sustaining the dynamic communication network underlying efficient Cas9 activation.

Our findings, together with our previous study on xCas9^12^, a variant of Cas9 that expands DNA recognition toward AAG and GAT PAM sequences, underscore a set of general biophysical rules that govern efficient PAM recognition in Cas9 variants. First, stabilization of the PAM-interacting domain is essential to ensure persistent base- and backbone-specific contacts, preventing local fluctuations from propagating into loss of global allosteric coupling. Inactive constructs that weaken the PI domain (e.g., through disruption of the D1135–K1107/S1109 network) fail to maintain PAM engagement despite retaining nominal recognition motifs. Second, long-range communication between the PI domain and the REC3 allosteric hub must be preserved, as demonstrated by our dynamic network analyses showing that efficient PAM binding propagates information to REC3 and onward to the HNH catalytic center. This coupling is diminished in inactive variants and reinforced in functional ones, highlighting PAM recognition as a distal allosteric switch. Third, entropic tuning of DNA binding, exemplified by the dynamic R1335 side chain in xCas9^12^, provides the flexibility required to accommodate diverse PAM sequences without incurring prohibitive entropic penalties. Together, these principles indicate that PAM recognition is not solely a matter of direct nucleotide contacts, but an emergent property of local stabilization, distal allosteric coupling, and entropic adaptability. Such rules provide a conceptual framework for engineering next-generation Cas9 variants with broadened PAM compatibility and enhanced activity.

## Conclusions

In summary, our computational study provides a unified mechanistic framework for understanding how PAM-site alterations reshape the allosteric circuitry of Cas9. By integrating MD simulations with graph-theoretical and centrality analyses, we show that efficient PAM recognition relies on three interdependent principles: stabilization of the PAM-binding domain, preservation of long-range allosteric communication with the REC3 hub, and entropic tuning of DNA engagement. Together, these features enable Cas9 to couple local PAM sensing with distal catalytic activation. Beyond clarifying the structural determinants of PAM specificity, our results establish general design rules that can guide the rational engineering of Cas9 variants with expanded PAM compatibility and enhanced genome-editing performance.

## Supporting Information

The Supporting Information is available free of charge at https://pubs.acs.org

## Author Contribution

FV and CP contributed equally to this work.

## Notes

The authors declare no competing interests.

## Acknowledgments

This material is based upon work supported by the NIH (Grant No. R01GM141329, to GP and Grant No. R01GM136815, to GP and GPL) and the NSF (Grant No. CHE-2144823, to GP and Grant No. MCB-2143760, to GPL). GP acknowledges support by the Alfred P. Sloan Foundation (Grant No. FG-2023-20431) and the Camille and Henry Dreyfus Foundation (Grant No. TC-24-063). The computational studies performed here were carried out using Expanse at the San Diego Supercomputing Centre through allocation MCB160059 and Bridges2 at the Pittsburgh Supercomputer Centre through allocation BIO230007 from the Advanced Cyberinfrastructure Coordination Ecosystem: Services & Support (ACCESS) program, which is supported by NSF support grants #2138259, #2138286, #2138307, #2137603, and #2138296.

